# Sticky enzymes: increased metabolic efficiency via substrate-dependent enzyme clustering

**DOI:** 10.1101/2024.11.05.622105

**Authors:** Alejandro Martínez-Calvo, Jello Zhou, Yaojun Zhang, Ned S. Wingreen

## Abstract

Coclustering of subsequent enzymes in a pathway can accelerate the processing of metabolic intermediates, with benefits including increased pathway fluxes, reduced toxicity, and sensitive branch-point regulation. While the optimal organization of such clusters has been explored theoretically, little is known about how to achieve such organization inside cells. Here we propose that phase-separating enzymes can self-organize into nearly-optimally sized and spaced clusters, provided that their “stickiness” is regulated by local substrate availability. In a nutshell, enzyme clusters only form when and where they are needed to process substrate. We study a mathematical model that implements this scheme for simple metabolic pathways, including all thermodynamic constraints. We find that pathway fluxes can be increased by 50 to 1000-fold and toxic metabolites can be decreased by 10 to 100-fold, at realistic enzyme densities. Finally, we discuss how enzyme “stickiness” could be allosterically regulated. This study presents a self-organization strategy that goes beyond current paradigms for natural and engineered enzyme clusters, and thus represents a motivating challenge to the fields of synthetic biology and metabolic engineering.

---

Colocalization of multiple enzymes in a pathway can enhance the rate of processing of intermediates, thereby improving the efficiency of cellular metabolism [1–15]. Consequently, considerable effort has been devoted to the experimental study of both natural and engineered enzyme clusters [16–21]. These intracellular metabolic assemblies, commonly referred to as *metabolons* or *metabolic condensates*, are known to create unique biochemical environments that facilitate substrate channeling and metabolite segregation at various steps in natural pathways. For example, purinosomes drive de novo purine biosynthesis [22–25] and glucosomes aid in glucose metabolism [3, 26–29]. The pyrenoid, a phase-separated organelle in algae, is another example in which colocalization of enzymes enhances photosynthetic carbon assimilation, accounting for nearly 30% of global carbon fixation [30–36]. In the context of synthetic biology and metabolic engineering, harnessing engineered enzyme clusters holds great promise for efficient biomanufacturing of fuels, materials, and pharmaceutical compounds, as well as for improving crop yields [37–39]. Additionally, detoxification and protection from toxic intermediates and reactive metabolites is one of the pillars of metabolic compartmentalization [2, 5, 8, 14, 38–44]. Since the efficiency of enzyme co-clusters depends on their spatio-temporal organization, it is important to understand, and ultimately control, how enzymes organize dynamically into favorable or even optimal configurations.

Key to the utility of both natural and engineered enzyme clusters is that they assemble and disassemble dynamically in response to the needs of the cell. For instance, the assembly/disassembly of purinosomes correlates with the cell’s demand for purine biosynthesis [24]. In the case of algae, pyrenoid formation is induced by CO_2_ availability and light [36]. Assembly/disassembly of these natural enzyme clusters is generally attributed to posttranslational protein modifications and requires operation of devoted regulatory systems [8, 16, 22, 23, 25, 36]. Assembly/disassembly of engineered clusters generally requires external control, e.g., via light exposure [8, 16, 17, 45–48]. While the mechanisms underpinning the basic assembly and disassembly of clusters have been explored in both natural and synthetic contexts, less attention has been paid to how cluster size and spatial organization are determined and how they can be controlled. Since metabolic efficiency can depend strongly on such spatial organization, it is natural to ask: how can cluster sizes and locations be regulated for greatest advantage? In particular, is it possible for clusters to dynamically self-organize, e.g., to achieve optimal pathway fluxes? While previous studies have proposed that enzyme clustering and self-organization can be achieved via diffusio- and chemo-phoresis [49–58]— enzymes migrating in response to a self-generated metabolite gradient [59]—it remains unexplored if such mechanisms can achieve dynamical self-organization into configurations that maximize the efficiency of metabolic pathways.

Here we make a simple proposal that functionally adaptive self-organization should depend on metabolite availability. In brief, enzymes should only cluster when and where they are needed to process metabolites. This makes sense intuitively, as the optimal size and spacing of clusters crucially depend on metabolite availability [2, 60]. To test this idea, we derive a minimal model that can achieve nearly optimal clusters when enzyme phase separation is regulated by metabolites through an allosteric site, distinct from the catalytic site. Allosteric regulation of enzymes is common in both natural and synthetic contexts [6, 61–68], and allostery operates as an equilibrium process, implying that no energy input is needed [69]. We find that allosterically-regulated enzyme clustering performs well over a range of different modeled conditions, e.g., when substrate is produced spatially uniformly in a two-step pathway or when substrate is produced at spatially localized sources of different strengths. Learning how to optimally control the spatial organization of enzyme clusters has implications for a broad range of applications in synthetic biology and metabolic engineering, as well as in other active and living systems whose constituents can self-organize in response to their environment.

## Results

### Model for substrate-dependent enzyme clustering via phase separation

Is it possible for enzymes to self-organize into nearly optimal clusters, thus maximizing the efficiency of a metabolic pathway? We first focus on a multi-step pathway. Assuming a particular cellular pathway has a fixed enzyme budget (i.e., total enzyme volume fraction), the optimal cluster size is the best compromise between the efficiency of consecutive steps of the pathway [2, 4, 60]. Aggregating all the enzymes into a few large clusters decreases the efficiency of the first step, since diffusion of a substrate produced throughout the cytoplasm to reach the nearest cluster becomes limiting. However, partitioning the enzymes into clusters that are too small decreases the efficiency of the second and any subsequent steps, as the metabolic intermediates produced inside the clusters can too quickly escape by diffusion before being processed. Theoretically, it is straightforward to calculate the optimally efficient cluster size, but in practice how can this optimal size be achieved inside cells?

Here, we propose a way that enzymes can exploit phase separation to self-organize into optimally sized clusters. To conceptually demonstrate this scheme, we consider a simple two-step metabolic pathway: Enzyme 1 converts substrate to intermediate, and Enzyme 2 converts intermediate to product (see Fig. **1**a). The key to self-organization is that Enzyme 1 has a sticky end that can drive phase separation, but the exposure of this sticky end is allosterically regulated by sub-strate binding to a site that is distinct from the catalytic site (see Fig. **1**b). Thus Enzyme 1 only forms clusters when and where its substrate is present. To colocalize Enzyme 2 into these clusters, we assume the two enzymes are connected by a short linker (alternatively, Enzyme 2 could colocalize with Enzyme 1 based on affinity). While in principle these ingredients could suffice to make enzymes form appropriately sized clusters when and where they are needed, there are many caveats to such self-organization, including thermodynamic constraints. Below we model this system in detail and show that such self-organized enzyme clusters are achievable with realistic parameters.

**Fig 1.**
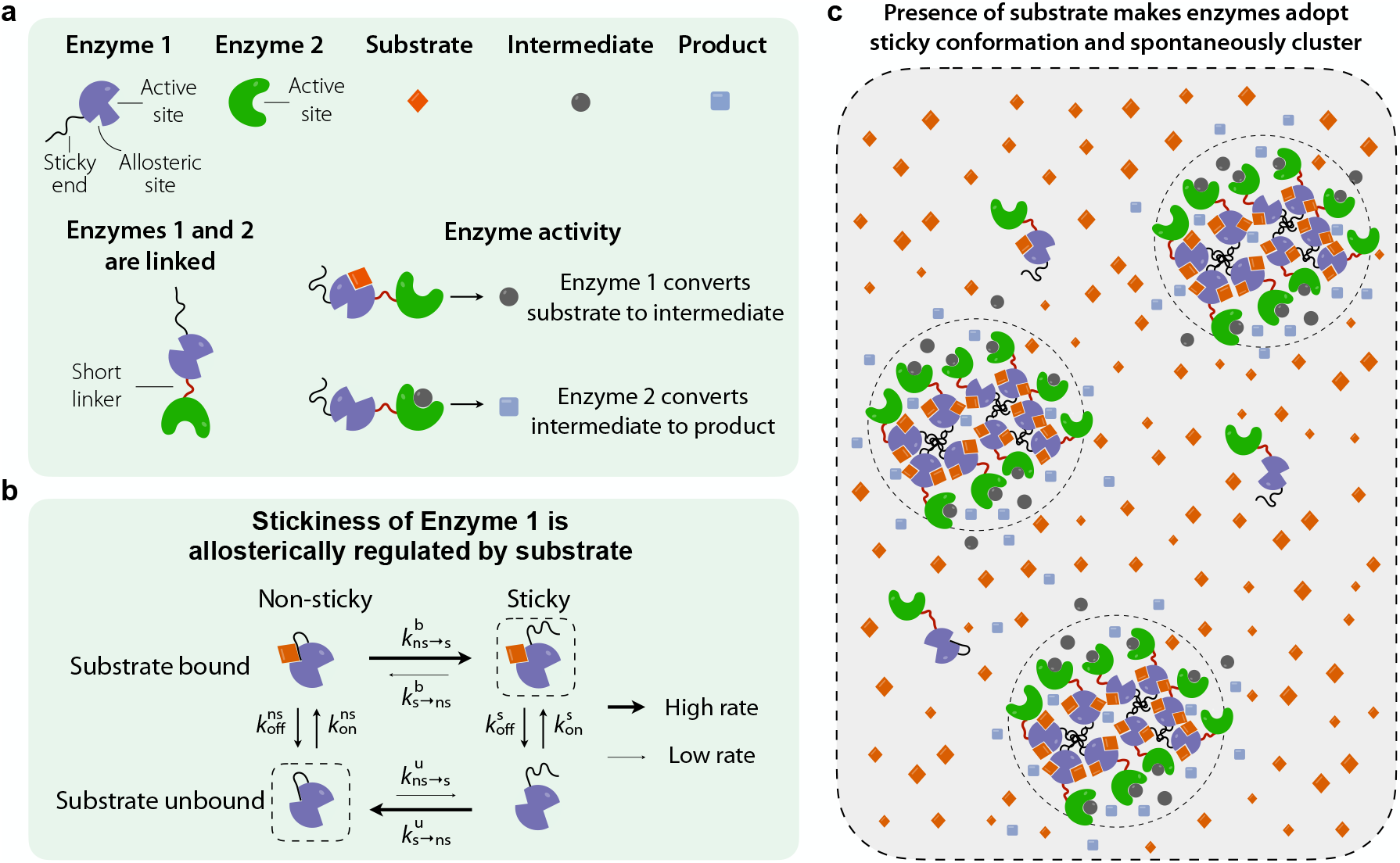
Substrate-dependent enzyme clustering via phase separation. **a**, Sketch of the enzymes and metabolites involved in a two-step metabolic pathway. Enzyme 1 and Enzyme 2 are connected by a short linker. Enzyme 1 converts substrate to intermediate, while Enzyme 2 converts intermediate to product. Enzyme 1 has a sticky end and can exist in either a sticky state or a non-sticky state, regulated by a non-catalytic allosteric site where substrate can bind. **b**, The rates of switching between these two states depend on the local substrate concentration, i.e., when substrate is bound to the allosteric site, the rate to become sticky is high, whereas when no substrate is bound the rate to become non-sticky is high. **c**, When substrate is present, enzyme clusters form spontaneously via phase separation of the sticky enzymes.

To model the metabolic pathway schematized in Fig. **1**a, we consider a suspension of enzyme complexes (each consisting of Enzyme 1 linked to Enzyme 2) diffusing in a solvent (the cytoplasm). We denote by *ϕ*_*i*_ the volume fraction of each of the three components of the mixture, where {*i* = s, ns, sol} denotes sticky enzyme complexes, non-sticky enzyme complexes, and solvent, respectively. We further assume that the substrate and intermediate are small molecules, whose concentrations *m*_1_(***r***, *t*) and *m*_2_(***r***, *t*) do not directly contribute to the volume fraction or other physical properties of the system.

### Enzyme free energies and chemical potentials

We model enzyme and solvent interactions via the following Flory-Huggins free-energy density [70–73]:

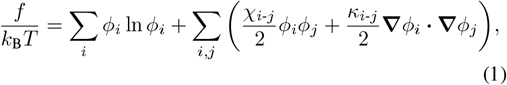

where *k*_B_ is the Boltzmann constant and *T* the absolute temperature. The first term in equation (1) is the entropy of mixing, which tends to keep the system well mixed. The second term represents the interaction energy between the components of the mixture, where the *χ*_*i*-*j*_ are Flory-Huggins interaction parameters. The third term reflects a surface energy that determines the width of the interface between phases. In equation (1) we have assumed that the molecular volume of enzymes and solvent molecules are the same, which is a reasonable approximation considering the crowded cytoplasm of the cell as solvent. In what follows, we also assume that non-sticky enzymes behave identically to solvent molecules, and therefore consider a single non-zero interaction parameter *χ*_s−s_ = *χ <* 0, stemming from the attractive interaction between the exposed sticky ends of Enzyme 1.

### Enzyme dynamics

To model the dynamics of enzymes, we consider the following kinetic equations:

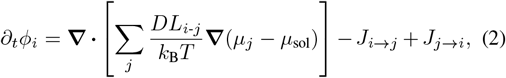

where *i, j* = {s, ns}, *D* is a constant diffusion coefficient, and the first term describes diffusive fluxes driven by gradients of chemical potential *µ*_*i*_, while *J*_*i*→*j*_ and *J*_*j*→*i*_ are the switching fluxes between sticky and non-sticky states. To derive equation (2) we have assumed that the diffusive fluxes are linearly proportional to gradients of the local chemical potentials *µ*, i.e., ***j***_*i*_ = −Σ_*j*_ *DL*_*i*-*j*_ ∇*µ*_*j*_*/*(*k*_B_*T*), where *L*_*i*-*j*_ is the matrix of mobility coefficients that satisfies the Onsager reciprocal relations, *L*_*i*-*j*_ = *L*_*j*-*i*_. Imposing an incompressibility condition Σ_*i*_ *ϕ*_*i*_ = 1 implies Σ_*i*_ ***j***_*i*_ = **0** and thus Σ_*j*_ *L*_*i*-*j*_ = 0, which yields equation (2) [74–76]. For simplicity, we assume that cross-terms are negligible, i.e., *L*_*i*-*j*_ = 0 for *i* ≠ *j*, so that gradients of the sticky enzyme chemical potential do not drive fluxes of non-sticky enzymes and vice versa, and *L*_*i*-*i*_ = *ϕ*_*i*_.

The chemical potential of sticky and non-sticky enzymes is obtained by minimizing the free energy of the system with respect to the conserved volume fractions, i.e., *µ*_*i*_ ≡ *δF/δϕ*_*i*_ = ∂*f/*∂*ϕ*_*i*_ − **∇** · ∂*f/*∂ **∇** *ϕ*_*i*_. Thus, the chemical potential differences driving the diffusive fluxes read (see Supplementary Information):

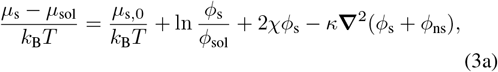

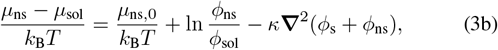

where *ϕ*_sol_ = 1 − *ϕ*_s_ − *ϕ*_ns_, and *µ*_s,0_ and *µ*_ns,0_ are bare chemical potentials of sticky and non-sticky states, respectively (see Supplementary Information for details on *κ*).

### Detailed balance of switching rates

Enzyme 1 can switch between sticky and non-sticky states, a conformational change regulated by a non-catalytic allosteric site (Fig. **1**a). Thus, Enzyme 1’s configuration depends on the local concentration of substrate *m*_1_(***r***, *t*). We assume that the binding-unbinding of substrate from the allosteric site and the switching between sticky and non-sticky configurations are equilibrium processes. To obtain the switching fluxes *J*_s→ns_ and *J*_ns→s_ between sticky and non-sticky configurations, we therefore impose detailed balance (i.e., no cyclic fluxes) on the four-state kinetic network sticky/non-sticky, substrate-bound/substrate-unbound (Fig. **1**b) [69]:

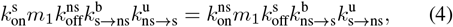

where 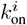 and 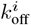 are the substrate binding and unbinding rates for sticky and non-sticky states, and 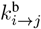 and 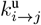 are the switching rates between sticky and non-sticky configurations for substrate-bound and unbound states, respectively. We define substrate dissociation constants of sticky and non-sticky states as 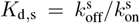 and 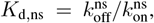, respectively. To obtain the total switching fluxes as functions of the local substrate concentration *m*_1_, it is useful to define the chemical potential difference Δ*µ/*(*k*_B_*T*) = (*µ*_s_ − *µ*_ns_)*/*(*k*_B_*T*) = Δ*µ*_0_*/*(*k*_B_*T*) + 2*χϕ*_s_ + ln(*ϕ*_s_*/ϕ*_ns_), where Δ*µ*_0_ is the bare chemical potential difference between sticky and non-sticky states, and the Boltzmann weight 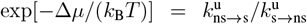, which implies 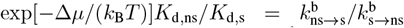 from equation (4). Eliminating 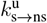 and 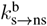 in favor of 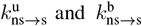, the total switching fluxes between sticky and non-sticky states can be written:

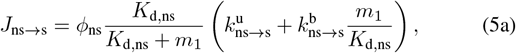

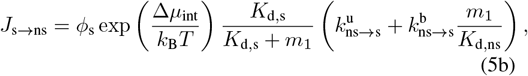

where Δ*µ*_int_*/*(*k*_B_*T*) = Δ*µ*_0_*/*(*k*_B_*T*) + 2*χϕ*_s_ is the interaction contribution to the total chemical potential difference Δ*µ*.

### Metabolites

For the two-step metabolic pathway, we consider the following reaction-diffusion equations for the substrate and intermediate,

Substrate:

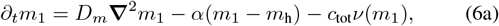

intermediate:

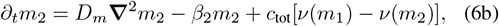

where *D*_*m*_ is a metabolite diffusion constant, assumed to be the same for both substrate and intermediate, *α* is the rate at which the substrate concentration relaxes everywhere to its homeostatic level *m*_h_, *c*_tot_ = *c*_max_(*ϕ*_s_ + *ϕ*_ns_) is the total concentration of sticky and non-sticky enzymes, with *c*_max_ the maximum possible molar concentration, and *β*_2_ is the intermediate decay rate. For simplicity, we assume that the processing of substrate by Enzyme 1 and intermediate by Enzyme 2 obey the same Michaelis-Menten kinetics, with maximal rate *k*_cat_ and dissociation constant *K*, i.e., *ν*(*m*) = *k*_cat_*m/*(*K* + *m*).

### Constraints on model parameters

For efficient processing of metabolites, enzymes must cluster only when and where they are needed. As a consequence: (i) Enzymes have to aggregate where the concentration of substrate is high. However, continued aggregation would lead to coarsening of clusters, finally yielding one big cluster containing all enzymes, which is far from an optimal organization as substrate diffusion into the cluster would be limiting. Thus, (ii) clusters must stop growing in size once substrate becomes substantially depleted in their interior, thereby yielding finite-size clusters that should be close to the optimal size. These two requirements imply certain mathematical constraints on the model parameters.

In our proposed model, (i) implies that the rate of switching from a non-sticky to a sticky configuration must be high where substrate is high, promoting aggregation. This is achieved by allosteric regulation: when substrate is bound to the allosteric site, it is likely that the sticky end will be free to interact with other sticky ends, thereby driving clustering (Fig. **1**b). To satisfy (ii), if a cluster is too big, such that substrate is depleted in its interior, sticky enzymes without substrate bound to their allosteric site must switch to the non-sticky state (i.e. sticky end not free to interact). Finally, to prevent continued coars-Sticky enzymes: increased metabolic efficiency via substrate-dependent enzyme clustering ening of enzyme clusters, non-sticky enzymes have to be able escape the cluster via diffusion faster than they switch back to the sticky state due to the increasing substrate concentrations toward the periphery of the cluster. Provided these constraints are satisfied, we expect that our model will produce a configuration of nearly optimally sized and spaced enzyme clusters. Quantitatively, what do the above constraints imply for the model parameters? Where substrate is locally high, i.e., *m*_1_ ≫ *K*_d,s_, *K*_d,ns_, the allosteric site is always occupied and the enzymes should be sticky, requiring *K*_d,ns_*/K*_d,s_ exp(−Δ*µ*_0_*/*(*k*_B_*T*) − 2*χϕ*_s_) ≫ 1, and this should also hold outside of clusters where *ϕ*_s_ ≃ 0. In practice we set *K*_d,ns_*/K*_d,s_ = 10^3^ − 10^4^, and also set *χ <* − 5 so that sticky enzymes phase separate even for *ϕ*_tot_ ≲ 0.1. In the opposite regime, where substrate is locally depleted, i.e., *m*_1_ ≪ *m*_h_ the homeostatic background substrate concentration, enzymes should be non-sticky, which implies exp(Δ*µ*_0_*/*(*k*_B_*T*) + 2*χϕ*_s_) ≫ 1. Inside substrate-depleted clusters, exp(2*χϕ*_s_) ≪ 1 as *ϕ*_s_ ≃ 1, reflecting the fact that the sticky state is energetically favored by interactions with the surrounding sticky enzymes, which restricts the allowed values of *χ* and Δ*µ*_0_. In practice we use |*χ*| = 5-6 and Δ*µ*_0_*/*(*k*_B_*T*) = 6-8, since a larger value of Δ*µ*_0_ would violate the first condition. These constraints make sense intuitively: where substrate is abundant, the sticky configuration should be energetically favored, whereas in the absence of substrate, the non-sticky configuration should be favored, even for an enzyme surrounded by sticky neighbors.

Finally, the time for a non-sticky enzyme to diffuse out of a cluster must be short compared to the time for it to switch back to a sticky state. This constraint implies 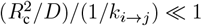, where *R*_c_ is the characteristic radius of an enzyme cluster. Assuming *R*_c_ ∼ 1 *µ*m and *D* = 1-100 *µ*m^2^, in practice we set *k*_*i*→*j*_ = 0.002 − 0.2 s^−1^.

### Self-organized enzyme clustering accelerates a two-step metabolic pathway

To determine whether our model can achieve a nearly optimal self-organization of enzymes, we perform full time-dependent numerical simulations of equations (2)–(6). We non-dimensionalize the equations using the substrate decay length 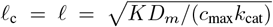 as a characteristic length scale, the enzyme diffusivity time scale *t*_c_ = ℓ^2^*/D* as a characteristic time scale, and the dissociation constant *K* as a characteristic scale of metabolite concentration (see Supplementary Information). We consider a cylindrical domain of dimensionless radius 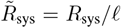 and length 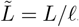, containing an initially 1:1 uniform mixture of sticky and non-sticky enzymes and uniform substrate and intermediate concentrations given by the uniform steady-state solution of equation (6). We solve the two-dimensional axisymmetric equations with periodic boundaries, having perturbed the system at initial time with small-amplitude white noise to trigger enzyme clustering.

We find that the enzymes self-organize into clusters of a fixed characteristic size (Fig. **2**a-c). To reach this state, enzyme complexes first become sticky because substrate concentration is uniformly high, and so spontaneously form dense clusters. These clusters coarsen, with some growing while others disappear, but eventually, the clusters reach a characteristic size. What keeps the clusters from continuing to coarsen? As a cluster grows, the concentration of substrate at its center decreases due to processing by the surrounding enzymes. Once the substrate concentration becomes low enough, enzymes at the center of the cluster begin to switch from sticky to non-sticky and diffuse out of the cluster. Once outside the cluster, these enzymes encounter higher levels of substrate and so switch back to being sticky. In our simulations, as clusters grow, eventually a steady-state cluster size is reached for which the rate of non-sticky enzymes leaving the cluster matches the rate of sticky enzymes joining the cluster.

In the two-step metabolic pathway described by equation (6), the advantage of forming clusters is to localize Enzyme 2 where the unstable intermediate is being produced by Enzyme 1. In the model we assume the two enzymes are connected, so consequently clusters of Enzyme 1 automatically contain an equally high density of Enzyme 2.

**Fig 2.**
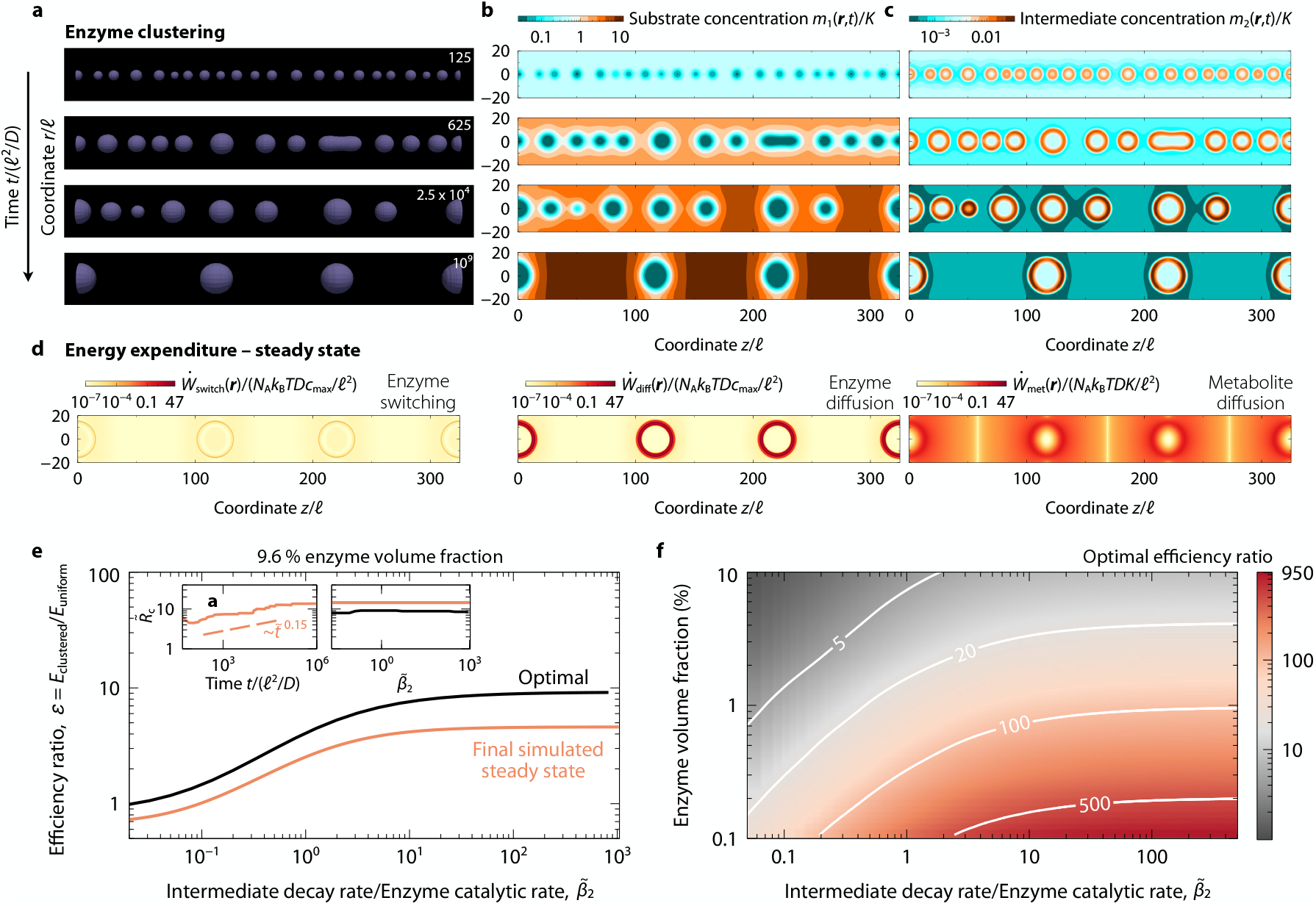
Substrate-dependent enzyme clustering accelerates a two-step metabolic pathway. **a**, Snapshots of enzyme clusters at dimensionless times *t/*(ℓ^2^*/D*) = 125, 625, 2.5 × 10^4^, and 10^9^, starting from a spatially uniform system with enzyme volume fraction *ϕ*_tot_ = 0.096, perturbed with small-amplitude white noise, in an axisymmetric cylindrical geometry. **b**, Dimensionless substrate concentration *m*_1_(***r***, *t*)*/K* at a slice taken at the mid-plane of the 3D clusters, at the same times as in panel **a**. When the substrate in the interior of the clusters becomes depleted by enzymatic activity, the enzymes become inactive and non-sticky, and diffuse away from the cluster, halting the coarsening of enzyme clusters. **c**, Dimensionless intermediate concentration *m*_2_(***r***, *t*)*/K* as in panel **b**. The values of dimensionless parameters are: 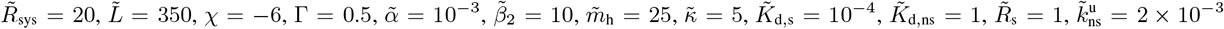, and Δ*µ*_0_*/*(*k*_B_*T*) = 8 (see Supplementary Information for details). **d**, Dimensionless energy dissipation rate 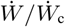 at steady state associated with enzyme switching (left), enzyme diffusion (middle), and metabolite diffusion (right). **e**, Ratio of efficiencies *ε* between clustered and uniform systems at final steady state, as a function of the ratio between intermediate decay rate and enzyme catalytic rate 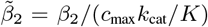, for dimensionless homeostatic concentration 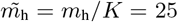 and homeostatic relaxation rate 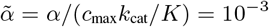 as in panels **a**–**c**. Solid curves correspond to numerical simulations at final steady state (orange) and optimal efficiency ratio (black). Left inset: characteristic cluster radius *R*_c_*/*ℓ as a function of time, computed from the simulation in panel **a**. Right inset: optimal and simulated cluster size 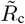 as a function of 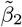. **f**, Color plot of the optimal clustered vs. uniform efficiency ratio as a function of total enzyme volume fraction *ϕ*_tot_ and dimensionless intermediate decay rate 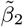.

What is the energy expenditure associated with the active self-organization of enzyme clusters? There are several distinct contributions to the energy dissipation: enzyme switching, enzyme diffusion, metabolite diffusion, and metabolite catalysis. While the last is arbitrary in our current model, since we assume reactions are strongly forward driven, the others are all calculable. Figure **2**d shows the spatial variation of the dimensionless dissipation (entropy production) rate 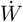 associated with enzyme switching (left), enzyme diffusion (middle), and metabolite diffusion (right), at steady state (see Supplementary Information). Although enzyme switching satisfies detailed balance, the dissipation rate is nonzero at steady state because the switching fluxes are influenced by the locally varying substrate concentration. Similarly, the gradients of sticky and non-sticky enzymes and of metabolites drive fluxes which also dissipate energy, even at steady state. All this energy ultimately comes from the catalysis of the metabolites. If this energy were not available, e.g., if we modeled the metabolic interactions as reversible and allowed them to reach true equilibrium, there would be no enzyme clustering.

To quantify how much pathway production is improved by enzyme clustering and to test if our model achieves a near-optimal enzyme configuration, we first compute the efficiency *E* of the pathway as the ratio between the total rate of product production and the maximal possible rate of production set by the maximal rate of substrate production:

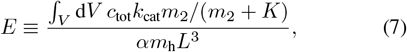

which is bounded between 0 ≤ *E* ≤ 1. To quantify the advantage of enzyme clustering, we then calculate the efficiency ratio between clustered and uniform configurations:

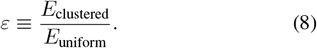

In Fig. **2**d we plot this efficiency ratio *ε* as a function of the (normalized) intermediate decay rate for the final steady-state cluster distribution (see Supplementary Information for details). We find that for a total enzyme volume fraction of 9.6%, self-organized enzyme clustering increases efficiency up to approximately 5-fold for high intermediate decay rates. How close is this to optimal? To estimate the optimal efficiency ratio, we consider that the overall enzyme distribution in a clustered system consists of a uniform pattern of well-separated dense clusters of identical radius *R*_c_ and uniform maximal volume fraction *ϕ*_tot_ = *ϕ*_s_ + *ϕ*_ns_ = 1. Thus, we only need to compute *ε* by solving equations (6) for one enzyme cluster and its surrounding volume *V*_c_ [2], and finding the optimal value of *R*_c_, and the corresponding *V*_c_, for a fixed volume fraction *ϕ*_tot_. As shown in Fig. **2**d, the self-organized clusters come within 60% of maximal efficiency over the entire range of decay rates tested (in the model, intermediate concentration does not affect clustering so the same self-organized solution applies for all intermediate decay rates.).

Importantly, the advantage of clustering increases at lower overall enzyme concentrations. While we are limited numerically to relatively high enzyme volume fractions ∼ 10%, typical values of certain enzymes in cells are in the range 0.1-5% [77–79]. For instance, in human cells (e.g., HeLa and HepG2 lines), purine and pyrimidine enzymes are ∼ 0.3% and ∼ 0.1-0.4% fraction of the total intracellular biomass, respectively [78, 79]. Glycolytic enzymes usually occupy a higher fraction of the cell’s biomass, ranging from ∼ 1-2% in human cells [78, 79], up to ∼ 7-10% in yeast cells [80, 81]. For this range of enzyme concentrations, optimal clustering can increase efficiency over the uniform case by ∼100-fold or more, as shown in Fig. **2**e. To quantitatively understand the magnitude of this effect, note that each intermediate molecule is either processed by Enzyme 2 or lost to decay. Since this processing rate is proportional to the local concentration of Enzyme 2 (as long as Enzyme 2 is unsaturated, i.e., *m*_2_*/K* ≪ 1, which is expected [2] and which we find to be the case; see Fig. **2**c), the increase of processing efficiency is approximately given by the ratio of Enzyme 2’s density in clusters to its density in the uniform case. For the choice of parameter values in Fig. **2**, *ϕ*_tot_ ∼ 0.1, thus *ε* ∼ 1*/ϕ*_tot_ ∼ 10, which is in good agreement with the increased metabolic efficiency obtained in Fig. **2**d. Note that this simple relation only applies when the intermediate decay rate is high enough that intermediates rarely escape from clusters by diffusion. This is the regime where clustering is most useful. Indeed, in the extreme limit of zero decay rate, all intermediates eventually get processed so clustering provides no advantage (Fig. **2**d and e).

### Spatial self-organization of enzymes around localized sub-strate sources

What if the production of substrate is not spatially uniform? For example, the metabolic building blocks for a pathway might be produced at particular spatial locations in the cell, e.g., at mitochrondria [22–25]. For the same reasons as above, enzyme clustering around localized sources of an unstable substrate should enhance the processing rate and improve yield. Here, we analyze whether our model of allosterically regulated phase-separating enzymes can generate appropriately sized clusters around localized substrate sources, as schematized in Fig. **3**a. We therefore now consider a one-step metabolic pathway in which substrate is converted to product by Enzyme 1, and compare the efficiency between clustered and uniform cases.

**Fig 3.**
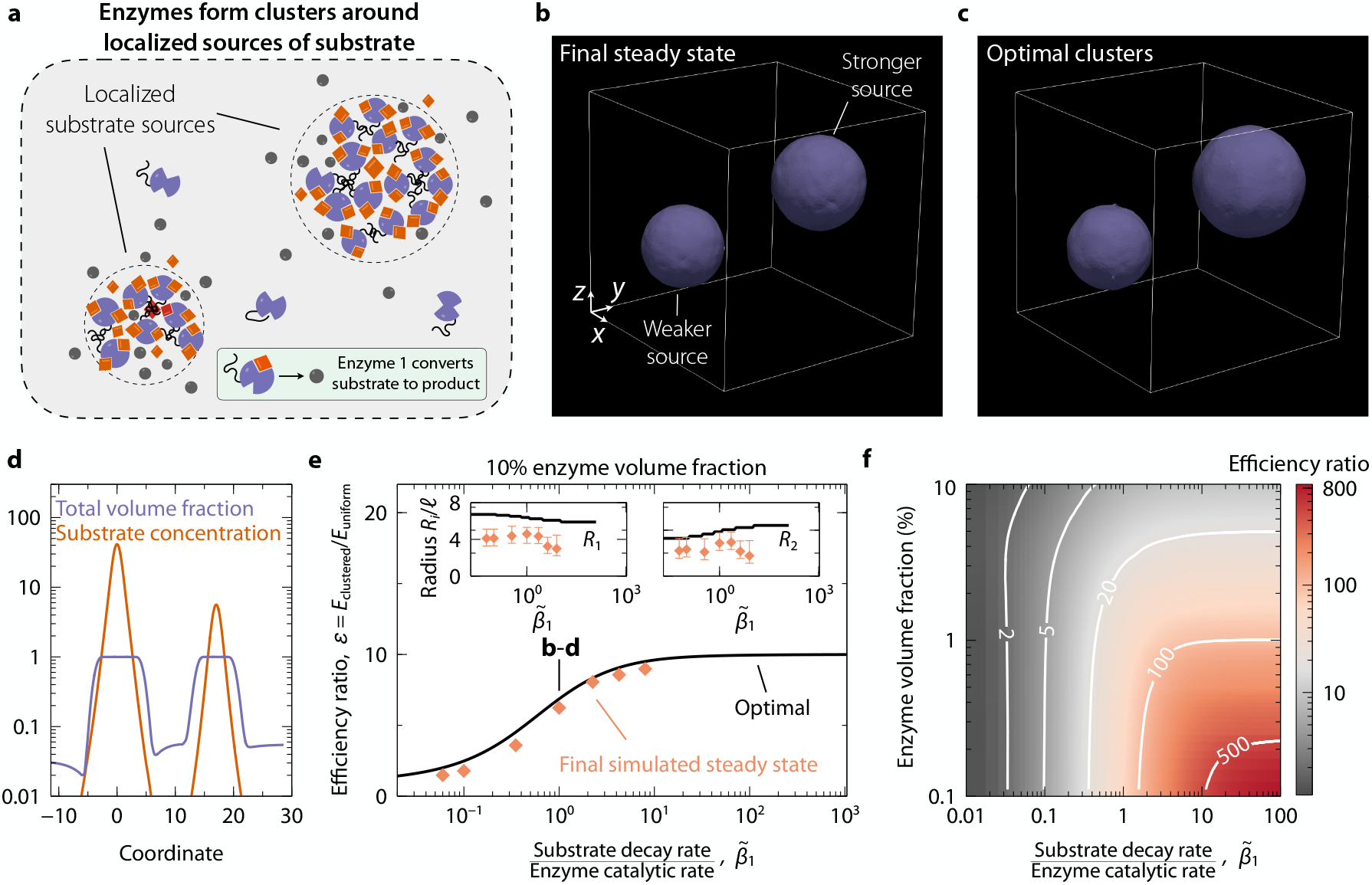
Nearly optimal efficiency is achieved by substrate-dependent enzyme clustering around spatially localized sources. **a**, Illustration of self-organized enzyme clusters around spatially localized substrate sources. **b, c**, Snapshots of enzyme clusters at the final simulated steady state (b), and from the optimal calculation (c) for two localized substrate sources, one 7 times as strong as the other. The total enzyme volume fraction is *ϕ*_tot_ = 0.1, and surfaces correspond to a local enzyme volume fraction of *ϕ*_tot_ = 0.1. The values of dimensionless parameters are: 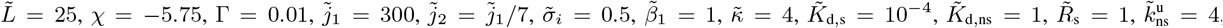, and Δ*µ*_0_*/*(*k*_B_*T*) = 5 (see Supplementary Information for details). **d**, Total volume fraction *ϕ*_tot_ and substrate concentration *m*_1_ along a line passing through the center of each cluster for the final simulated steady state. **e**, Ratio of efficiencies between clustered and uniform systems, for both simulated (orange symbols) and optimal (black solid curve) cases, as a function of ratio between substrate decay rate and enzyme catalytic rate 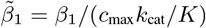. Inset: Simulated (orange symbols) and optimal (black solid curve) cluster radii *R*_*i*_*/*ℓ as functions of 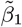. Symbols represent a radius at which *ϕ*_tot_ = 0.5, with lower and upper error bars corresponding to *ϕ*_tot_ = 0.9 and 0.1, respectively. **f**, Optimal efficiency ratio between clustered and uniform systems as a function of total enzyme volume fraction and substrate decay rate 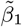.

To account for localized substrate sources, equation (6a) becomes:

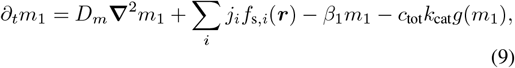

where *f*_s,*i*_(***r***) is a narrow Gaussian function centered at location ***r***_s,*i*_ (see Supplementary Information), and *j*_*i*_ is the source strength. The substrate is assumed to be unstable with a decay rate *β*_1_. The equations for enzymes are the same, i.e., equations (2) and (5). We initialize the simulations with zero substrate, *m*_1_(***r***, *t* = 0) = 0, and a uniform volume fraction of only non-sticky enzymes, i.e., *ϕ*_ns_(***r***, *t* = 0) = *ϕ*_tot_ and *ϕ*_s_(***r***, *t* = 0) = 0. In this case, we compute the efficiency as follows:

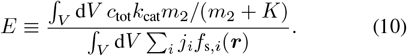

We performed full time-dependent numerical simulations of equations (2), (5) and (9) for a case of 10% enzyme volume fraction and two localized sources of substrate, one 3 times as strong as the other (see Supplementary Information). Figure **3**b shows the final simulated steady state in which clusters have formed around the localized sources. Figure **3**d displays the total enzyme volume fraction *ϕ*_tot_(***r***) and sub-strate concentration *m*_1_(***r***) along a line passing through the centers of both clusters at steady state (panel b), showing that only a negligible amount of substrate escapes the enzyme clusters. Figure **3**c displays the optimal configuration of clusters, i.e., the enzyme distribution into two maximally dense clusters centered around two localized sources that maximizes the efficiency of the pathway. We find good agreement between the optimal and simulated cluster sizes. Indeed, Fig. **3**e shows that the simulated clustered/uniform efficiency ratio reaches at least 90% of the optimal value over the range of substrate decay rates tested. In particular, for a 10% enzyme volume fraction and a fixed value of the system size normalized by the metabolite decay length, 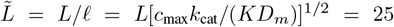, we find that enzyme clustering increases efficiency up to ∼10-fold when the dimensionless substrate decay rate 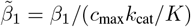 is sufficiently high. As in the uniform substrate case, the gain in efficiency increases with lower volume fraction, as depicted in Fig. **3**f, which shows the efficiency ratio as a function of enzyme volume fraction and dimensionless substrate decay rate 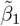. The insets to Fig. **3**e show the radii of the two clusters as a function of 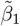. Our model yields somewhat smaller clusters than the optimal calculation because the smaller clusters already achieve nearly maximal processing of substrate, so there is no advantage to adding more enzymes to the clusters.

### Toxic metabolites

Many metabolic reactions, such as purine biosynthesis and galactose processing, can produce intermediates or end products that are toxic to cells, e.g., resulting in damage or the inhibition of other reactions [2, 5, 8, 14, 38–44]. Moreover, even intermediates that are nontoxic contribute to the overall osmolarity of the cell and therefore have a cost in terms of osmotic stress and consequent loss of function [82]. Thus, it is interesting to explore whether enzymes can self-organize into clusters that optimally reduce the total amount of a toxic metabolite. For simplicity we again consider a one-step pathway (though this could be part of a longer pathway) where the substrate is stable but toxic. To this end, we perform the same numerical simulations of equations (2), (5) and (9), for a non-decaying “toxic” substrate.

Figure **4**a shows the total spatially averaged substrate concentration as a function of time, i.e. ∫_*V*_ d*V m*_1_(***r***, *t*), for both clustered and uniform cases, revealing that enzyme clustering reduces the total amount of toxin substantially. Here, the enzyme volume fraction is 8%, resulting in a ∼ 5-fold decrease of total toxin when enzymes cluster around the source. Figure **4**b displays the total volume fraction of enzymes *ϕ*_tot_(***r***) and toxic substrate concentration *m*_1_(***r***) along a cross-section through the center of the source, at the final simulated steady state (panel a, iii), for both clustered and uniform cases. We find that enzyme clustering accelerates substrate processing and thus reduces both the maximum and overall toxic substrate concentration.

To highlight the relevance of enzyme volume fraction, here we also compute the ratio of toxin between clustered and uniform systems for a single source and an enzyme cluster of uniform volume fraction *ϕ*_tot_ = *ϕ*_s_ + *ϕ*_ns_ = 1, centered at the source. To this end, we solve equation (9) at steady state for different enzyme volume fractions *ϕ*_tot_. Figure **4**c shows this ratio as a function of *ϕ*_tot_ for different values of the system size normalized by the toxin decay length, 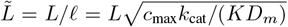. Interestingly, we find that there is an optimal amount of enzymes, i.e., an optimal value of *ϕ*_tot_, that minimizes the clustered/uniform toxin ratio for a fixed value 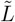, and that therefore maximizes the toxin reduction when enzymes are clustered. This result makes sense intuitively: for a fixed metabolite diffusion constant *D*_*m*_ and enzyme catalytic rate *c*_max_*k*_cat_*/K*, there is an optimal cluster radius below which a substantial amount of substrate escapes by diffusion before getting processed, and above which very little substrate reaches the periphery of the cluster.

**Fig 4.**
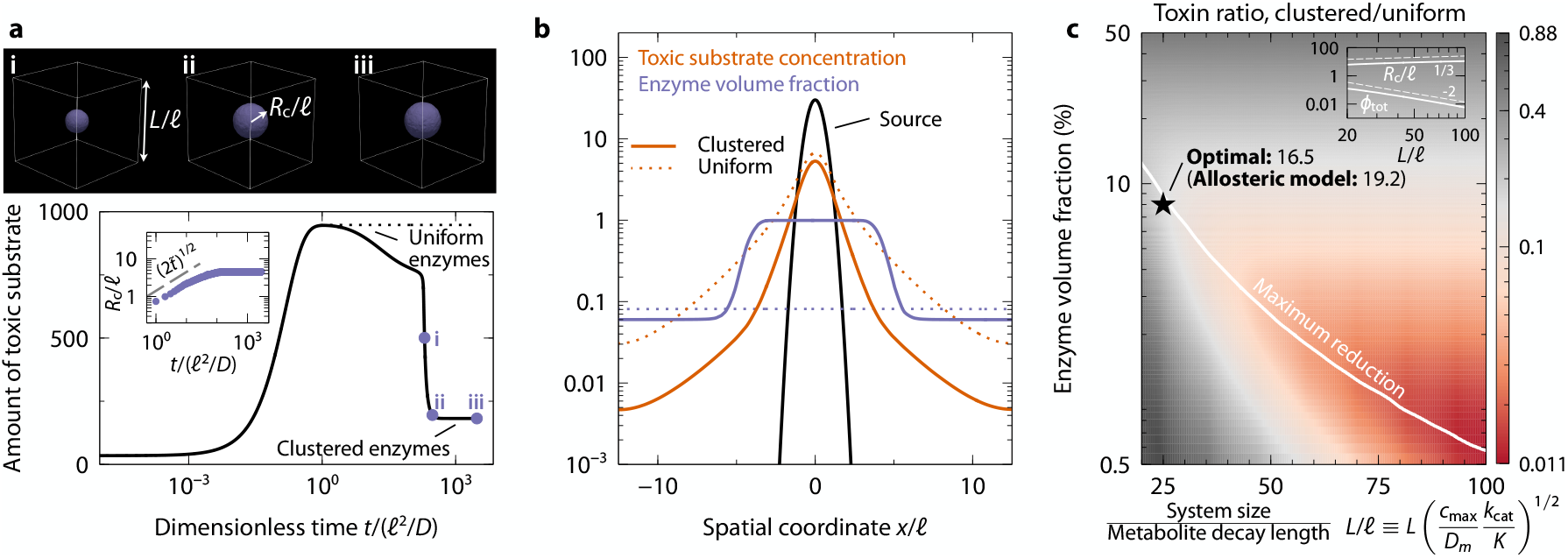
Substrate-dependent enzyme clustering can reduce toxic burden. **a**, Amount of toxic substrate as a function of time for both uniform (dashed curve) and clustered (solid curve) cases. Inset: Dimensionless cluster radius *R*_c_*/*ℓ as a function of time *t/*(ℓ^2^*/D*). The snapshots i-iii display the aggregation of enzymes around a source of toxic substrate at different times indicated with blue dots along the curve in the main panel. **b**, Toxic substrate concentration and total enzyme volume fraction along a cross-section through the center of the source, for the case of uniform enzyme distribution (dashed curves) and spontaneous enzyme clustering via phase separation (solid curves). Here, 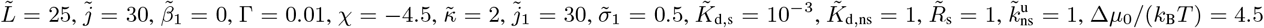 and *ϕ*_tot_ = 0.08. **c**, Optimal toxin ratio between clustered and uniform systems as a function of enzyme volume fraction *ϕ*_tot_ and dimensionless system size 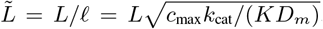. The dot corresponds to the steady-state ratio of panel a, and the curve corresponds to the maximum toxin reduction. The inset shows *ϕ*_tot_ and *R*_c_*/*ℓ as a function of *L/*ℓ along the maximum reduction curve.

Our model yields a near optimal toxin reduction (star in Fig. **4**c). The main difference from the optimal result stems from not having all the available enzymes in the cluster. As for the case of localized sources in Fig. **3**, it is the enzymes at the periphery of clusters that encounter low substrate levels and consequently transition to a non-sticky state. This self-organized configuration could prove advantageous in alternative scenarios, as these unclustered enzymes are available elsewhere where their functionality might be required.

### Dynamics of cluster formation

How long does it take our modeled enzyme clusters to self-organize? In the case of a spatially uniform substrate concentration, clusters emerge and grow initially via spinodal decomposition until coarsening is arrested and clusters reach a stable equilibrium size. We find that the average cluster size, computed as the inverse of the first moment of the structure factor, scales as 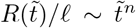 with *n* 0.1 0.2, where 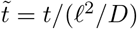 (left inset to Fig. **2**d; see Supplementary Information). Thus, enzyme switching to non-sticky states makes cluster growth slower than conventional Ostwald ripening for which *n* = 1*/*3 as dictated by the Lifshitz-Slyozov growth law [83]. Both slower cluster growth and arrested coarsening are common features of reactive and active mixtures, where the presence of nonequilibrium reactions and nonreciprocal interactions between system components suppresses long-wavelength modes [58, 72, 73, 84–90].

In the case of localized substrate sources, we find that clusters initially grow in a diffusive fashion, i.e., *R*_c_(*t*) *t* ∼ ^1*/*2^, before arresting. This power-law growth can be estimated by computing the diffusion-limited flux of enzymes onto a sticky sphere, i.e., assuming that any non-sticky enzymes switch to a sticky state prior to contacting the sphere, leading to *J* = 4*πρDR*_c_, where *ρ* is the density of enzymes far from the sphere. Therefore, the sphere volume will grow as 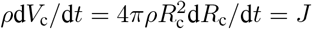, which yields *R*_c_ = (2*Dt*)^1*/*2^ (see inset to Fig. **4**a; note that *R*_c_(*t* = 0) depends on an initial transient period before the onset of diffusion-limited growth, accounting for the *y*-axis offset with respect to the data).

## Discussion

Clustering of multiple enzymes in a pathway can improve the efficiency of cellular metabolism by accelerating the processing of metabolic intermediates. While efficiency gains depend strongly on the spatiotemporal organization of such clusters [2, 60], little is known about how in practice to achieve optimally sized and spaced clusters. In this work, we propose that enzymes can exploit phase separation to self-organize into nearly optimal configurations, provided that their attractive interactions are regulated by local substrate availability. Thus, enzymes cluster only when and where they are needed to process metabolites. To test this idea, we developed a minimal continuum model in which enzymes can switch between sticky and non-sticky states, regulated by a non-catalytic allosteric site. We find that such enzymes can robustly self-organize into nearly optimally sized and spaced clusters under a variety of conditions, e.g., when substrate is produced uniformly in space or instead at spatially localized sources. Our results also revealed that this clustering strategy is nearly optimal in minimizing the amount of stable metabolic intermediates, thus preventing damage outside clusters if these are toxic or reactions with other compounds in the bulk, which might result in futile cycles or other toxic by-products.

*What is necessary for enzymes to optimally self-organize?* Key to proper enzyme self-organization is that where substrate is depleted, enzymes should *not* accumulate, so the conversion rate to the non-sticky state should be high. Conversely, where substrate concentration is high, the switching rate to the sticky state should be high. In addition, enzyme switching rates need to be slow enough that non-sticky enzymes inside clusters can diffuse away before switching back to the sticky state. These requirements impose constraints on the allosteric parameters, as well as on the enzyme’s maximal switching rates. Moreover, since allostery is an equilibrium process, there are constraints associated with satisfying detailed balance among the four enzyme states shown in Fig. **1**b. Despite equilibrium switching, the system’s dissipation rate at steady state is nonzero because of the spatially varying metabolite concentrations. The energy source for enzyme clustering ultimately comes from metabolite catalysis. Allowing for an additional energy input to control switching would remove some of the constraints on switching rates, but would require more complex regulation and additional energy expenditure. However, this is not necessary–––indeed we have shown that the allosteric model can function for realistic parameter values. Moreover, engineering enzyme regulation via allosteric switches, usually referred to as ‘modular allostery’, is an established method, inspired by native systems. [6, 61–68, 91]. *Alternative approaches and future extensions of our work*. An alternative approach based on equilibrium processes and close to our minimal scheme is for allostery to drive the formation of dimers or larger oligomers that would phase separate, even when monomers would not, e.g., ligand-induced dimerization [92, 93]. This is a very common mechanism for regulating phase separation in native [94–97] and optogenetic systems [8, 17, 18, 45, 48]. Another plausible equilibrium self-organization strategy is for an allosteric site to be regulated by the final product of the pathway, in such a way that when product levels are low, enzyme clustering is favored. Nonetheless, many native phase-separating systems operate out of equilibrium and require energy sources, such as kinases and phosphatases or RNA production or helicase activity [98, 99]. Indeed, an alternative to allosteric regulation would be to couple switching of an enzyme to its catalytic activity. While as noted above such a nonequilibrium mechanism would presumably be substantially more complicated at the protein chemistry level and therefore more difficult to engineer, extending our model and ideas to out-of-equilibrium regulation of enzyme phase separation is a relevant direction for future work.

In this work, we have only explored short metabolic pathways. Would this self-organization strategy work for longer pathways? For the case of unstable intermediates, previous work has shown that the efficiency gains of clustering are multiplicative with each step of the pathway, and thus can be very large [2]. Formation of such multi-enzyme clusters could be achieved via linkers connecting different enzymes, as in the current study, or by means of enzyme-enzyme affinity interactions, or combinations of the two. Furthermore, while we have explored the energy expenditure associated with short pathways, how does energy dissipation scale with multiple steps when enzymes self-organize into clusters? What self-organization strategies minimize energy expenditure while still optimizing yield? These are intriguing future research directions.

Another important property of biochemical systems is component stoichiometry, which can also impact the formation, internal organization, and properties of intracellular condensates [100–105]. Here, for simplicity and ease of engineering, we have considered the simplest possible scenario of two enzymes with 1:1 stoichiometry. This can be achieved by physically connecting the enzymes with a short linker as we propose, or by engineering the two enzymes as separate proteins that form heterodimers. However, what if the optimal stoichiometry is not 1:1 or needs to adapt? Fixed non 1:1 stoichiometries could be achieved via more complex linkages or by engineering downstream enzymes to be clients of Enzyme 1 clusters, requiring engineering Enzyme 1 to have multiple binding sites for the other enzymes. An intriguing possibility is that adaptive self-organization of stoichiometry could be achieved if downstream enzymes are also regulated allosterically by their corresponding substrates, thereby clustering when and where these enzymes are needed.

Slow and/or arrested coarsening of native and synthetic biomolecular condensates have been reported previously, e.g., experimentally in light-activated droplets [17, 48, 106], and theoretically in reactive and active mixtures, in the presence of nonequilibrium reactions and nonreciprocal interactions between system components [58, 72, 73, 84–90]. Moreover, recent theoretical works have considered enzyme aggregation via diffusiophoresis, in which enzymes can migrate along chemical gradients [49–57, 107]. It is not yet clear whether versions of these mechanisms could lead to optimal enzyme organization, as in most cases they lead to continued coarsening of clusters. Exploring whether the same principles of substrate-dependent clustering can also be achieved by some of these distinct mechanisms is a promising direction for future research.

## Conclusions

This work addresses a question that is central to cellular metabolism and metabolic compartmentalization: how can enzymes self-organize into configurations that maximize yield? Here, we presented a novel self-organization strategy based on substrate-dependent phase separation that goes beyond the current paradigms for native and synthetic enzyme assemblies. We hope our results will inspire and guide original experiments in metabolic engineering. Moreover, our findings are not restricted to metabolism, as the spatiotemporal regulation of biomolecular condensates is crucial to many other cellular processes, including transcription [98, 108–111], cell signaling [100, 112], and stress responses [113–115]. More generally, our work may also relate to a broad range of systems at the supracellular level, whose constituents can dynamically self-organize, regulated by their environment and metabolic activity. These include synthetic matter [116, 117] and living systems such as microbial communities [118–122].

A.M.-C. acknowledges support from the Princeton Center for Theoretical Science, the Center for the Physics of Biological Function, and the Human Frontier Science Program through the grant LT000035/2021-C. N.S.W. acknowledges support from the NSF through the Center for the Physics of Biological Function PHY-1734030 and the NIH through grant R01 GM140032. We also thank the Princeton Biomolecular Condensate Program for funding support. We thank Zemer Gitai, Trevor GrandPre, and Qiwei Yu for thoughtful discussions.

## SUPPLEMENTARY INFORMATION

### Free energy and chemical potentials

The free-energy density of the system is given by equation 1 of the Main Text, whose full expression in terms of *ϕ*_s_, *ϕ*_ns_, and *ϕ*_sol_ reads:

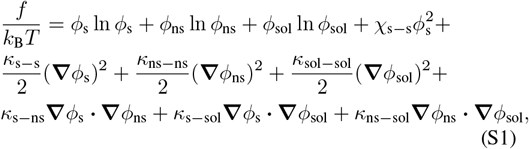

where for simplicity we have assumed that non-sticky enzymes have the same interaction parameters as solvent, and therefore consider a single non-zero interaction parameter *χ*_s−s_ = *χ <* 0, stemming from the attractive interaction between the exposed sticky ends of Enzyme 1. We minimize *f* with respect to *ϕ*_s_, *ϕ*_ns_, and *ϕ*_sol_ to obtain the chemical potentials of sticky and non-sticky enzyme complexes, and solvent, respectively:

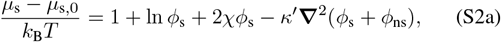

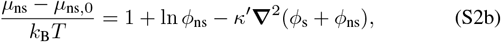

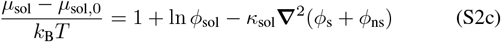

where *µ*_*i*,0_ are bare chemical potentials, and for simplicity we have assumed *κ*^*′*^ = *κ*_s-s_ − *κ*_s-sol_ = *κ*_s-ns_ − *κ*_s-sol_ = *κ*_s-ns_ − *κ*_ns-sol_ = *κ*_ns-ns_ − *κ*_ns-sol_, and *κ*_sol_ = *κ*_s-sol_ − *κ*_sol-sol_ = *κ*_ns-sol_ − *κ*_sol-sol_. the co nservation equations ∂_*t*_*ϕ*_*i*_ + **∇** · ***j***_*i*_ = 0, which yield

### Incompressibility constraint

To ensure incompressibility, i.e., Σ_*i*_ *ϕ*_*i*_ = 1, the diffusive fluxes ***j***_*i*_ need to satisfy some constraints [74–76]. First, we impose incompressibility on the conservation equations *∂*_*t*_*ϕ*_*t*_ + **∇** · *j*_*i*_ = 0 which yield **∇** ·(Σ_*i*_ ***j***_*i*_) = 0. Next, we Σ_*i*_ *j*_*i*_ = 0. assume that the fluxes are linearly proportional to gradients of chemical potential, i.e. *j*_*i*_ = *D/*(*k*_B_*T*) Σ_*j*_ *L*_*i*-*j*_ **∇** *µ*_*j*_, which for the case of our three-component mixture read:

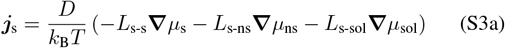

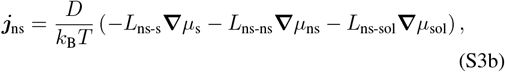

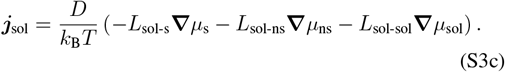

Imposing the incompressibility constraint Σ_*i*_ *j*_*i*_ yields the following relations between Onsager coefficients, Σ_*j*_ *L*_*i*-*j*_ = 0, that allow us to eliminate all coefficients related to the solvent: *L*_s-sol_ = −(*L*_s-s_ + *L*_s-ns_), *L*_ns-sol_ = −(*L*_s-ns_ + *L*_ns-ns_), and *L*_sol-sol_ = *L*_s-s_ + *L*_ns-ns_ + 2*L*_s-ns_ where we have already imposed the Onsager reciprocal relations, *L*_*i*-*j*_ = *L*_*j*-*i*_. Thus, the fluxes for sticky and non-sticky enzymes now read:

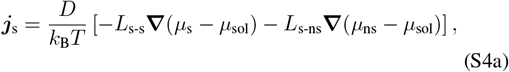

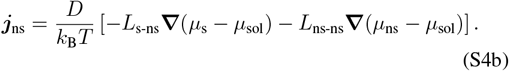

To eliminate *ϕ*_sol_ from the solvent chemical potential *µ*_sol_, we impose Σ_*i*_ *ϕ*_*i*_ = 1, which yields two closed equations for sticky *ϕ*_s_ and non-sticky *ϕ*_ns_ enzyme volume fractions. For simplicity, we further assume that cross-diffusive terms are negligible, which implies *L*_s-ns_ = 0, and *L*_*i*-*i*_ = *ϕ*_*i*_, yielding equation (2) of the Main Text. We expect that our results are, in principle, not significantly altered by this choice. The chemical potential differences between sticky/non-sticky states and the solvent driving the diffusive fluxes read:

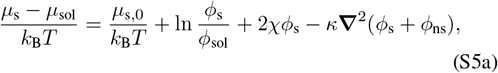

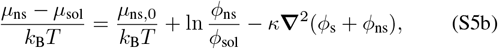

where *κ* = *κ*^*′*^ − *κ*_sol_, *ϕ*_sol_ = 1 − *ϕ*_s_ − *ϕ*_ns_, and *µ*_s,0_ and *µ*_ns,0_ are bare chemical potentials of sticky and non-sticky states, respectively. For simplicity and without loss of generality we have assumed that *µ*_sol,0_ = 0.

### Derivation of allosteric switching rates

We consider that the conformational change between the sticky and non-sticky state of enzymes is regulated by a non-catalytic allosteric site where substrate can bind. Thus, this site exhibits 4 states as depicted in the 4-state reaction network shown in Fig. **1**b of the Main Text. The binding and unbinding rates are denoted by 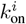 and 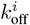, respectively, where *i* = {s, ns}, and the switching rates between sticky and non-sticky states are denoted by 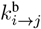 and 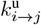, when substrate is bound and unbound, respectively. Since allostery is an equilibrium process, this network must satisfy detailed balance, i.e., no cyclic fluxes. We assume that the binding-unbinding of substrate to the allosteric site is much faster than local changes in the concentration of substrate, which allows us to write the binding rates proportional to the substrate concentration. Thus, detailed balance implies:

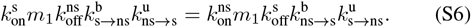

Hence, the kinetics simplify to just equilibrium state occupancies that depend on the dissociation constants, 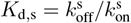 and 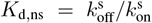, along with an offset energy Δ*µ* = *µ*_s_ − *µ*_ns_ between the sticky and non-sticky states in the absence of substrate. Thus, the probabilities of the four states read:

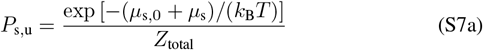

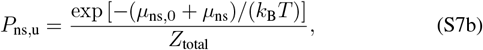

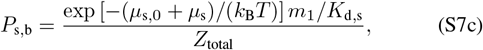

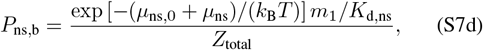

where *Z*_total_ is the partition function of the system that ensures that probabilities of states sum up to one, and which reads:

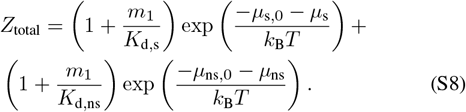

We compute the total rates of switching from the sticky state to the non-sticky state and vice versa as follows:

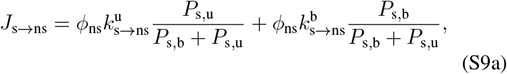

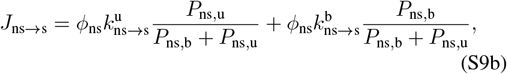

which are the equations, together with the conservation equations of metabolites (equation (6) of the Main Text), we solve numerically to obtain the results of the Main Text.

### Non-dimensionalization and dimensionless parameters

To reduce the number of parameters of the model and identify the relevant ones, we non-dimensionalize equations (2)– where we have multiplied both switching fluxes by *ϕ*_ns_ without loss of generality. This choice results in a switching flux from non-sticky to sticky and sticky to non-sticky proportional to *ϕ*_ns_ and *ϕ*_s_, respectively. From detailed balance we know that 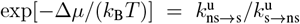 and 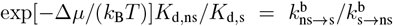. Eliminating 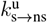 and 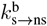 in favor of 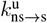 and 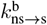, yield the following expressions for the total switching fluxes between sticky and non-sticky states:

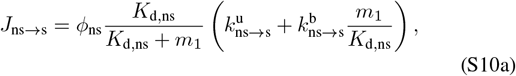

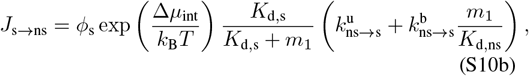

where Δ*µ*_int_*/*(*k*_B_*T*) = Δ*µ*_0_*/*(*k*_B_*T*) + 2*χϕ*_s_ is the interaction contribution to the total chemical potential difference Δ*µ*, and Δ*µ*_0_ = *µ*_s,0_ − *µ*_ns,0_ is the bare chemical potential difference between sticky and non-sticky states.

### Full equations of enzyme dynamics

Introducing the expression for the chemical potentials [equation (3) of the Main Text] into the conservation equations for sticky and non-sticky enzymes [equation (2) of the Main Text] result in the following coupled equations:

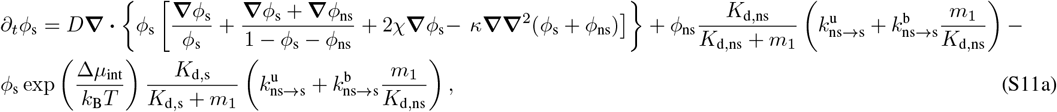

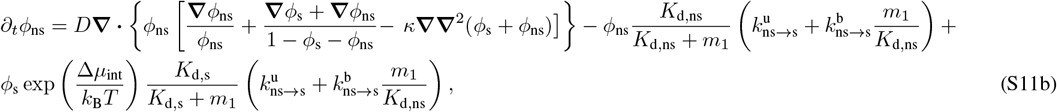

(6) of the Main Text. To this end, we consider the substrate penetration length 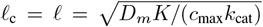 as characteristic length scale, i.e. the length in which substrate diffusion and catalysis are of the same order, and which sets how much substrate is able to penetrate into enzyme clusters. As characteristic time scale, we consider the time it takes enzyme to diffuse the characteristic length scale ℓ_c_, i.e., *t*_c_ = ℓ^2^*/D* = *D*_*m*_*K/*(*c*_max_*k*_cat_*D*). As for the substrate and intermediate concentration scales, we consider the Michaelis-Menten constant *c*_c_ = *K*. All dimensionless variables, except volume fractions, are denoted with tilde. The governing dimensionless parameters are:

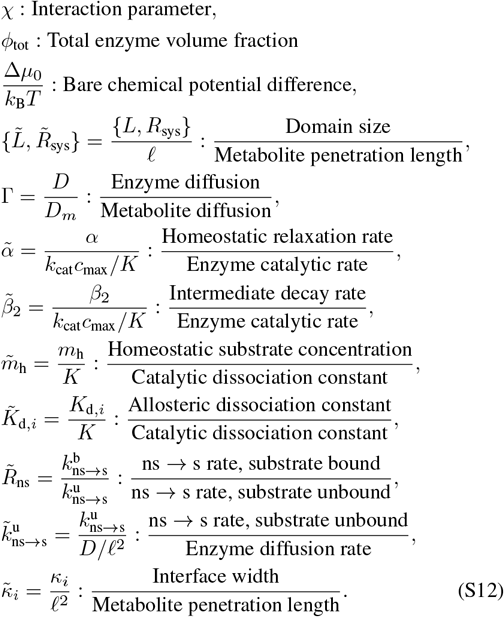

### Localized substrate sources

To model the localized substrate sources, we consider the following Gaussian functions:

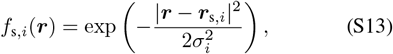

where ***r***_s,*i*_ is the position vector of the localized substrate source and *σ*_*i*_ determines the decay length of the metabolite source. Hence, the equation for the substrate (equation (9) of the Main Text) reads in dimensionless form:

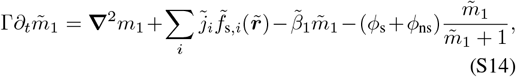

where 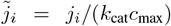 compares the rate of substrate production of each source with the enzyme catalytic rate, 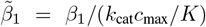 compares the substrate decay rate with the enzyme catalytic rate, and 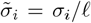 compares the decay length of the metabolite source with the metabolite penetration length.

### Characteristic enzyme cluster size

In the the case where substrate is produced uniformly in space, we compute the characteristic cluster size over time from the structure factor [76, 84, 135, 136]:

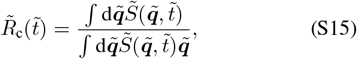

where 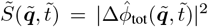 is the structure factor of the enzyme amplitude profile Δ*ϕ*_tot_ = *ϕ*_s_ + *ϕ*_ns_ − *ϕ*_tot,0_, and 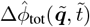 is its Fourier transform.

### Energy expenditure of enzyme switching and diffusion

We compute the dissipation (entropy production) rates associated with enzyme switching between sticky and non-sticky states 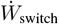, enzyme diffusion, 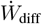, and metabolite diffusion 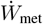 which read:

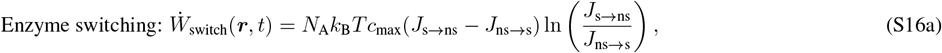

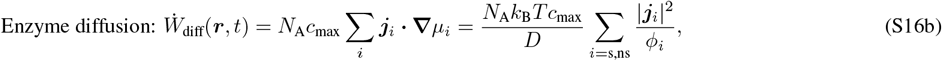

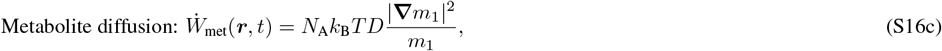

where *N*_A_ is Avogadro’s number.

### Values of model parameters

Table S1 displays the values of the model parameters used in the simulations of the Main Text.

**Table S1.** Values of relevant model parameters.

| Physical parameters | Definition | Range of values | Reference |
| --- | --- | --- | --- |
| $D_m$ | Metabolite diffusivity | 100–1000 $\mu\text{m}^2 \text{s}^{-1}$ | Refs. [2, 123, 124] |
| $D$ | Enzyme diffusivity | 0.1–100 $\mu\text{m}^2 \text{s}^{-1}$ | Refs. [125–128] |
| $c_{\text{max}}$ | Maximum enzyme concentration | $10^{-2}$–25 mM | Ref. [2, 129] |
| $k_{\text{cat}}/K$ | Enzyme catalytic rate | $10 - 5 \times 10^4 \text{ (mM s)}^{-1}$ | Refs. [2, 130–132] |
| $K$ | Catalytic dissociation constant | 1–1000 $\mu\text{M}$ | Refs. [2, 132, 133] |
| $m_h$ | Homeostatic metabolite concentration | 0.01 – 5 mM | Ref. [134] |
| $\alpha$ | Substrate homeostatic relaxation rate | $0.1 \text{ s}^{-1}$ | Ref. [2] |
| $\beta_2$ | Intermediate decay rate | $10^{-4}$–$10 \text{ s}^{-1}$ | Ref. [2] |
| $k_{i \rightarrow j}^u = k_{i \rightarrow j}^b$ | Switching rates | 0.002 – 0.2 $\text{s}^{-1}$ | |
| **Characteristic scales** |  |  |  |
| $\ell_c = \sqrt{D_m/(c_{\text{max}}k_{\text{cat}}/K)}$ | Characteristic length scale | 10 nm – 100 $\mu\text{m}$ | |
| $t_c = \ell_c^2/D$ | Characteristic time scale | $10^{-6} \text{ s}$ – 30 h | |
| **Dimensionless parameters** |  |  |  |
| $\Gamma = D/D_m$ | Enzyme diffusion/Metabolite diffusion | $10^{-4}$–1 | |
| $\tilde{m}_h = m_h/K$ | Homeostatic concentration/Dissociation constant | 0.01–5000 | |
| $\tilde{\alpha} = \alpha/(k_{\text{cat}}c_{\text{max}}/K)$ | Homeostatic relaxation rate/Enzyme catalytic rate | $10^{-7}$–1 | |
| $\tilde{\beta}_2 = \beta_2/(k_{\text{cat}}c_{\text{max}}/K)$ | Intermediate rate/Enzyme catalytic rate | $10^{-11}$–$10^2$ | |
| $\tilde{k}_{\text{ns} \rightarrow \text{s}}^u = k_{\text{ns} \rightarrow \text{s}}^u/(D/\ell_c^2)$ | Enzyme switching rate/Enzyme diffusion rate | $10^{-5}$–$10^6$ | |

## References

[1] R. J. Conrado, J. D. Varner, and M. P. DeLisa, Curr. Opin. Biotechnol. 19, 492 (2008).

[2] M. Castellana, M. Wilson, Y. Xu, P. Joshi, I. Cristea, J. Rabinowitz, Z. Gitai, and N. Wingreen, Nat. Biotech. 32, 1011 (2014).

[3] S. Jang, J. Nelson, E. Bend, L. Rodríguez-Laureano, F. Tueros, L. Cartagenova, K. Underwood, E. Jorgensen, and D. Colón-Ramos, Neuron 90, 278 (2016).

[4] F. Hinzpeter, U. Gerland, and F. Tostevin, Biophys. J. 112, 767 (2017).

[5] S. Banani, H. Lee, A. Hyman, and M. Rosen, Nat. Rev. Mol. Cell Biol. 18, 285 (2017).

[6] M. Prouteau and R. Loewith, Biomol. 8, 160 (2018).

[7] S. Tsitkov and H. Hess, ACS Catal. 9, 2432 (2019).

[8] E. M. Zhao, N. Suek, M. Z. Wilson, E. Dine, N. L. Pannucci, Z. Gitai, J. L. Avalos, and J. E. Toettcher, Nat. Chem. Biol. 15, 589 (2019).

[9] M. Liu, S. He, L. Cheng, J. Qu, and J. Xia, Biomacromolecules 21, 2391 (2020).

[10] A. Lyon, W. Peeples, and M. Rosen, Nat. Rev. Mol. Cell Biol. 22, 215 (2021).

[11] B. O’Flynn and T. Mittag, Curr. Opin. Cell Biol. 69, 70 (2021).

[12] V. Pareek, Z. Sha, J. He, N. Wingreen, and S. Benkovic, Mol. Cell 81, 3775 (2021).

[13] S. Kondrat, U. Krauss, and E. von Lieres, Curr. Res. Chem. Biol., 100031 (2022).

[14] L. Bar-Peled and N. Kory, Nat. Metab. 4, 1232 (2022).

[15] J. Breger, J. Vranish, E. Oh, M. Stewart, K. Susumu, G. Lasarte-Aragonés, G. Ellis, S. Walper, S. Díaz, S. Hooe, et al., Nat. Commun. 14, 1757 (2023).

[16] Y. Shin and C. Brangwynne, Science 357, eaaf4382 (2017).

[17] Y. Shin, J. Berry, N. Pannucci, M. P. Haataja, J. E. Toettcher, and C. P. Brangwynne, Cell 168, 159 (2017).

[18] D. Bracha, M. T. Walls, and C. P. Brangwynne, Nat. Biotech. 37, 1435 (2019).

[19] Y. Dai, L. You, and A. Chilkoti, Nat. Rev. Bioeng. 1, 466 (2023).

[20] Y. Dai, M. Farag, D. Lee, X. Zeng, K. Kim, H.-I. Son, X. Guo, J. Su, N. Peterson, J. Mohammed, et al., Nat. Chem. Biol. 19, 518 (2023).

[21] M. Walls, K. Xu, C. Brangwynne, and J. Avalos, bioRxiv, 2023 (2023).

[22] S. An, R. Kumar, E. Sheets, and S. Benkovic, Science 320, 103 (2008).

[23] H. Zhao, J. French, Y. Fang, and S. Benkovic, Chem. Commun. 49, 4444 (2013).

[24] C. Chan, H. Zhao, R. Pugh, A. Pedley, J. French, S. Jones, X. Zhuang, H. Jinnah, T. Huang, and S. Benkovic, Proc. Natl. Acad. Sci. U.S.A. 112, 1368 (2015).

[25] A. Pedley and S. Benkovic, Trends Biochem. Sci. 42, 141 (2017).

[26] C. Kohnhorst, M. Kyoung, M. Jeon, D. Schmitt, E. Kennedy, J. Ramirez, S. Bracey, B. Luu, S. Russell, and S. An, J. Biol. Chem. 292, 9191 (2017).

[27] M. Jin, G. Fuller, T. Han, Y. Yao, A. Alessi, M. Freeberg, N. Roach, J. Moresco, A. Karnovsky, M. Baba, et al., Cell Rep. 20, 895 (2017).

[28] S. An, M. Jeon, E. Kennedy, and M. Kyoung, in Methods in enzymology, Vol. 628 (Elsevier, 2019) pp. 1–17.

[29] M. Jeon, D. Schmitt, M. Kyoung, and S. An, bioRxiv, 2022 (2022).

[30] L. Mackinder, M. Meyer, T. Mettler-Altmann, V. Chen, M. Mitchell, O. Caspari, E. Freeman Rosenzweig, L. Pallesen, G. Reeves, A. Itakura, et al., Proc. Natl. Acad. Sci. U.S.A. 113, 5958 (2016).

[31] E. S. F. Rosenzweig, B. Xu, L. K. Cuellar, A. Martinez-Sanchez, M. Schaffer, M. Strauss, H. N. Cartwright, P. Ronceray, J. M. Plitzko, F. Förster, et al., Cell 171, 148 (2017).

[32] T. Wunder, S. Cheng, S.-K. Lai, H.-Y. Li, and O. Mueller-Cajar, Nat. Commun. 9, 5076 (2018).

[33] H. Wang, X. Yan, H. Aigner, A. Bracher, N. Nguyen, W. Hee, B. Long, G. Price, F. Hartl, and M. Hayer-Hartl, Nature 566, 131 (2019).

[34] J. H. Hennacy and M. C. Jonikas, Annu. Rev. Plant Biol. 71, 461 (2020).

[35] C. Fei, A. T. Wilson, N. M. Mangan, N. S. Wingreen, and M. C. Jonikas, Nat. Plants 8, 583 (2022).

[36] S. He, V. L. Crans, and M. C. Jonikas, Plant Cell, koad157 (2023).

[37] J. Nielsen and J. Keasling, Cell 164, 1185 (2016).

[38] S. Hammer and J. Avalos, Nat. Chem. Biol. 13, 823 (2017).

[39] N. Kulagina, S. Besseau, N. Papon, and V. Courdavault, Front. Bioeng. Biotechnol. 9, 659431 (2021).

[40] H. Hürlimann, B. Laloo, B. Simon-Kayser, C. Saint-Marc, F. Coulpier, S. Lemoine, B. Daignan-Fornier, and B. Pinson, J. Biol. Chem. 286, 30994 (2011).

[41] A. Bar-Even, A. Flamholz, E. Noor, and R. Milo, Nat. Chem. Biol. 8, 509 (2012).

[42] K. McGary, J. Slot, and A. Rokas, Proc. Natl. Acad. Sci. U.S.A. 110, 11481 (2013).

[43] H. Huttanus and X. Feng, Biotechnol. J. 12, 1700052 (2017).

[44] C. Kerfeld, C. Aussignargues, J. Zarzycki, F. Cai, and M. Sutter, Nat. Rev. Microbiol. 16, 277 (2018).

[45] D. Bracha, M. T. Walls, M.-T. Wei, L. Zhu, M. Kurian, J. L. Avalos, J. E. Toettcher, and C. P. Brangwynne, Cell 175, 1467 (2018).

[46] L. Zhu, T. M. Richardson, L. Wacheul, M.-T. Wei, M. Feric, G. Whitney, D. L. J. Lafontaine, and C. P. Brangwynne, Proc. Natl. Acad. Sci. U.S.A. 116, 17330 (2019).

[47] M.-T. Wei, Y.-C. Chang, S. F. Shimobayashi, Y. Shin, A. R. Strom, and C. P. Brangwynne, Nat. Cell Biol. 22, 1187 (2020).

[48] D. S. W. Lee, C.-H. Choi, D. W. Sanders, L. Beckers, J. A. Riback, C. P. Brangwynne, and N. S. Wingreen, Nat. Phys. 19, 586 (2023).

[49] J. Agudo-Canalejo, P. Illien, and R. Golestanian, Nano Lett. 18, 2711 (2018).

[50] J. Agudo-Canalejo, T. Adeleke-Larodo, P. Illien, and R. Golestanian, Acc. Chem. Res. 51, 2365 (2018).

[51] J. Agudo-Canalejo and R. Golestanian, Phys. Rev. Lett. 123, 018101 (2019).

[52] A. Testa, M. Dindo, A. Rebane, B. Nasouri, R. Style, R. Golestanian, E. Dufresne, and P. Laurino, Nat. Commun. 12, 6293 (2021).

[53] M. Cotton, R. Golestanian, and J. Agudo-Canalejo, Phys. Rev. Lett. 129, 158101 (2022).

[54] V. Ouazan-Reboul, J. Agudo-Canalejo, and R. Golestanian, Eur. Phys. J. E 44, 1 (2021).

[55] V. Ouazan-Reboul, J. Agudo-Canalejo, and R. Golestanian, arXiv preprint 2303.09832 (2023).

[56] V. Ouazan-Reboul, R. Golestanian, and J. Agudo-Canalejo, arXiv preprint 2305.05472 (2023).

[57] V. Ouazan-Reboul, R. Golestanian, and J. Agudo-Canalejo, arXiv preprint 2304.09925 (2023).

[58] L. Demarchi, A. Goychuk, I. Maryshev, and E. Frey, Phys. Rev. Lett. 130, 128401 (2023).

[59] S. Sengupta, K. Dey, H. Muddana, T. Tabouillot, M. Ibele, P. Butler, and A. Sen, J. Am. Chem. Soc. 135, 1406 (2013).

[60] A. Buchner, F. Tostevin, and U. Gerland, Phys. Rev. Lett. 110, 208104 (2013).

[61] F. Nord and C. Werkman, Advances in enzymology and related areas of molecular biology (John Wiley & Sons, 1941).

[62] T. Traut, Crit. Rev. Biochem. Mol. Biol. 29, 125 (1994).

[63] J. Smith, E. Zaluzec, J.-P. Wery, L. Niu, R. Switzer, H. Zalkin, and Y. Satow, Science 264, 1427 (1994).

[64] A. Mattevi, M. Bolognesi, and G. Valentini, FEBS Lett. 389, 15 (1996).

[65] W. Lim, Curr. Opin. Struct. Biol. 12, 61 (2002).

[66] I. Bahar, C. Chennubhotla, and D. Tobi, Curr Opin Struct Biol. 17, 633 (2007).

[67] P. Mehrabi, C. Di Pietrantonio, T. Kim, A. Sljoka, K. Taverner, C. Ing, N. Kruglyak, R. Pomes, E. Pai, and R. Prosser, J. Am. Chem. Soc. 141, 11540 (2019).

[68] R. Alberstein, A. Guo, and T. Kortemme, Curr. Opin. Struct. Biol. 72, 71 (2022).

[69] T. Einav, L. Mazutis, and R. Phillips, J. Phys. Chem. B 120, 6021 (2016).

[70] P. J. Flory, J. Chem. Phys. 10, 51 (1942).

[71] M. L. Huggins, J. Am. Chem. Soc. 64, 1712 (1942).

[72] J. Berry, C. Brangwynne, and M. Haataja, Rep. Prog. Phys. 81, 046601 (2018).

[73] C. Weber, D. Zwicker, F. Jülicher, and C. Lee, Rep. Prog. Phys. 82, 064601 (2019).

[74] E. J. Kramer, P. Green, and C. J. Palmstrøm, Polymer 25, 473 (1984).

[75] K. W. Kehr, K. Binder, and S. M. Reulein, Phys. Rev. B 39, 4891 (1989).

[76] S. Mao, D. Kuldinow, M. P. Haataja, and A. Košmrlj, Soft Matter 15, 1297 (2019).

[77] W. Duckworth, R. Bennett, and F. Hamel, Endocr. Rev. 19, 608 (1998).

[78] N. Nagaraj, J. Wisniewski, T. Geiger, J. Cox, M. Kircher, J. Kelso, S. Pääbo, and M. Mann, Mol. Syst. Biol. 7, 548 (2011).

[79] T. Geiger, A. Wehner, C. Schaab, J. Cox, and M. Mann, Mol. Cell. Proteom. 11 (2012).

[80] L. De Godoy, J. Olsen, J. Cox, M. Nielsen, N. Hubner, F. Fröhlich, T. Walther, and M. Mann, Nature 455, 1251 (2008).

[81] N. Nagaraj, N. Kulak, J. Cox, N. Neuhauser, K. Mayr, O. Hoerning, O. Vorm, and M. Mann, Mol. Cell. Proteom. 11 (2012).

[82] D. Kültz, Annu. Rev. Physiol. 67, 225 (2005).

[83] I. Lifshitz and V. Slyozov, J. Phys. Chem. Solids 19, 35 (1961).

[84] S. C. Glotzer, E. A. Di Marzio, and M. Muthukumar, Phys. Rev. Lett. 74, 2034 (1995).

[85] M. Bazant, Farad. Disc. 199, 423 (2017).

[86] Y. Li and M. Cates, J. Stat. Mech. Theory Exp. 2020, 053206 (2020).

[87] Z. You, A. Baskaran, and M. Marchetti, Proc. Natl. Acad. Sci. U.S.A. 117, 19767 (2020).

[88] S. Saha, J. Agudo-Canalejo, and R. Golestanian, Phys. Rev. X 10, 041009 (2020).

[89] J. Kirschbaum and D. Zwicker, J. R. Soc. Interface 18, 20210255 (2021).

[90] F. Brauns, H. Weyer, J. Halatek, J. Yoon, and E. Frey, Phys. Rev. Lett. 126, 104101 (2021).

[91] J. Dueber, B. Yeh, K. Chak, and W. Lim, Science 301, 1904 (2003).

[92] J. Klemm, S. Schreiber, and G. Crabtree, Annu. Rev. Immunol. 16, 569 (1998).

[93] E. Mack, R. Perez-Castillejos, Z. Suo, and G. Whitesides, Anal. Chem. 80, 5550 (2008).

[94] M. Feric, N. Vaidya, T. Harmon, D. Mitrea, L. Zhu, T. Richardson, R. Kriwacki, R. Pappu, and C. Brangwynne, Cell 165, 1686 (2016).

[95] D. Mitrea, J. Cika, C. Guy, D. Ban, P. Banerjee, C. Stanley, A. Nourse, A. Deniz, and R. Kriwacki, eLife 5, e13571 (2016).

[96] S. Aoki, T. Lynch, S. Crittenden, C. Bingman, M. Wickens, and J. Kimble, Nat. Commun. 12, 996 (2021).

[97] H. Tourrière, K. Chebli, L. Zekri, B. Courselaud, J. Blanchard, E. Bertrand, and J. Tazi, J. Cell Biol. 222 (2023).

[98] J. Henninger, O. Oksuz, K. Shrinivas, I. Sagi, G. LeRoy, M. Zheng, J. Andrews, A. Zamudio, C. Lazaris, N. Hannett, et al., Cell 184, 207 (2021).

[99] D. Overwijn and M. Hondele, Trends Biochem. Sci. 48, 244 (2023).

[100] L. B. Case, X. Zhang, J. A. Ditlev, and M. K. Rosen, Science 363, 1093 (2019).

[101] J.-M. Choi, F. Dar, and R. V. Pappu, PLoS Comput. Biol. 15, e1007028 (2019).

[102] D. W. Sanders, N. Kedersha, D. S. W. Lee, A. R. Strom, V. Drake, J. A. Riback, D. Bracha, J. M. Eeftens, A. Iwanicki, A. Wang, et al., Cell 181, 306 (2020).

[103] I. Alshareedah, G. M. Thurston, and P. R. Banerjee, Biophysical journal 120, 1161 (2021).

[104] Y. Zhang, B. Xu, B. G. Weiner, Y. Meir, and N. S. Wingreen, eLife 10, e62403 (2021).

[105] A. G. T. Pyo, Y. Zhang, and N. S. Wingreen, Iscience 25 (2022).

[106] D. Lee, N. Wingreen, and C. Brangwynne, Nat. Phys. 17, 531 (2021).

[107] X. Zhao, H. Palacci, V. Yadav, M. Spiering, M. Gilson, P. Butler, H. Hess, S. Benkovic, and A. Sen, Nat. Chem. 10, 311 (2018).

[108] J. Berry, S. Weber, N. Vaidya, M. Haataja, and C. Brangwynne, Proc Natl Acad Sci U.S.A. 112, E5237 (2015).

[109] B. Sabari, A. Dall’Agnese, A. Boija, I. Klein, E. Coffey, K. Shrinivas, B. Abraham, N. Hannett, A. Zamudio, J. Manteiga, et al., Science 361, eaar3958 (2018).

[110] A. Boija, I. Klein, B. Sabari, A. Dall’Agnese, E. Coffey, A. Zamudio, C. Li, K. Shrinivas, J. Manteiga, N. Hannett, et al., Cell 175, 1842 (2018).

[111] M. Du, S. Stitzinger, J.-H. Spille, W.-K. Cho, C. Lee, M. Hijaz, A. Quintana, and I. Cissé, Cell 187, 331 (2024).

[112] X. Su, J. Ditlev, E. Hui, W. Xing, S. Banjade, J. Okrut, D. King, J. Taunton, M. Rosen, and R. Vale, Science 352, 595 (2016).

[113] J. Riback, C. Katanski, J. Kear-Scott, E. Pilipenko, A. Rojek, T. Sosnick, and D. Drummond, Cell 168, 1028 (2017).

[114] C. Iserman, C. Altamirano, C. Jegers, U. Friedrich, T. Zarin, A. Fritsch, M. Mittasch, A. Domingues, L. Hersemann, M. Jahnel, et al., Cell 181, 818 (2020).

[115] S. Yasuda, H. Tsuchiya, A. Kaiho, Q. Guo, K. Ikeuchi, A. Endo, N. Arai, F. Ohtake, S. Murata, T. Inada, et al., Nature 578, 296 (2020).

[116] A. Dinsmore, M. Hsu, M. Nikolaides, M. Marquez, A. Bausch, and D. Weitz, Science 298, 1006 (2002).

[117] S. Ceron, G. Gardi, K. Petersen, and M. Sitti, Proc. Natl. Acad. Sci. U.S.A. 120, e2221913120 (2023).

[118] E. Ben-Jacob, I. Cohen, and H. Levine, Adv. Phys. 49, 395 (2000).

[119] Y. Guo, M. Tikhonov, and M. Brenner, Proc. Natl. Acad. Sci. U.S.A. 115, 3593 (2018).

[120] J. A. Schwartzman, A. Ebrahimi, G. Chadwick, Y. Sato, B. R. K. Roller, V. J. Orphan, and O. X. Cordero, Curr. Biol. 32, 3059 (2022).

[121] A. Dal Co, M. Ackermann, and S. van Vliet, Cell Syst. 14, 98 (2023).

[122] G. D’Souza, A. Ebrahimi, A. Stubbusch, M. Daniels, J. Keegstra, R. Stocker, O. Cordero, and M. Ackermann, ISME J. 17, 703 (2023).

[123] A. Garcia-Perez, E. Lopez-Beltran, P. Kluner, J. Luque, P. Ballesteros, and S. Cerdan, Arch. Biochem. Biophys. 362, 329 (1999).

[124] W. Stein and T. Litman, Channels, carriers, and pumps: an introduction to membrane transport (Elsevier, 2014).

[125] C. Riedel, R. Gabizon, C. Wilson, K. Hamadani, K. Tsekouras, S. Marqusee, S. Pressé, and C. Bustamante, Nature 517, 227 (2015).

[126] Y. Zhang and H. Hess, ACS Cent. Sci. 5, 939 (2019).

[127] M. Xu, J. Ross, L. Valdez, and A. Sen, Phys. Rev. Lett. 123, 128101 (2019).

[128] G. He, T. GrandPre, H. Wilson, Y. Zhang, M. Jonikas, N. Wingreen, and Q. Wang, Commun. Biol. 6, 19 (2023).

[129] M. Choe, T. Einav, R. Phillips, and D. Titov, bioRxiv, 2022 (2022).

[130] L. Kuo, A. Miller, S. Lee, and C. Kozuma, Biochem. 27, 8823 (1988).

[131] F. Guillou, M. Liao, A. Garcia-Espana, and C. Lusty, Biochem. 31, 1656 (1992).

[132] W. Yu, K. Jin, D. Wang, N. Wang, Y. Li, Y. Liu, J. Li, G. Du, X. Lv, J. Chen, et al., Nat. Commun. 15, 7989 (2024).

[133] J. Berg, J. Tymoczko, and L. Stryer, Biochemistry (Macmillan, 2007).

[134] C. Chassagnole, N. Noisommit-Rizzi, J. Schmid, K. Mauch, and M. Reuss, Biotechnol. Bioeng. 79, 53 (2002).

[135] H. Furukawa, Phys. Rev. E 61, 1423 (2000).

[136] V. M. Kendon, M. E. Cates, I. Pagonabarraga, J. C. Desplat, and P. Bladon, J. Fluid Mech. 440, 147 (2001).

